# An interplay between extracellular signalling and the dynamics of the exit from pluripotency drives cell fate decisions in mouse ES cells

**DOI:** 10.1101/000653

**Authors:** David A. Turner, Jamie Trott, Penelope Hayward, Pau Rué, Alfonso Martinez Arias

## Abstract

Embryonic Stem cells derived from the epiblast tissue of the mammalian blastocyst, retain the capability to differentiate into any adult cell type and are able to self-renew indefinitely under appropriate culture conditions. Despite the large amount of knowledge that we have accumulated to date about the regulation and control of self-renewal, efficient directed differentiation into specific tissues remains elusive. In this work, we have analyzed in a systematic manner the interaction between the dynamics of loss of pluripotency and Activin/Nodal, BMP4 and Wnt signalling in fate assignment during the early stages of differentiation of mouse ES cells in culture. During the initial period of differentiation cells exit from pluripotency and enter an Epi-like state. Following this transient stage, and under the influence of Activin/Nodal and BMP signalling, cells face a fate choice between differentiating into neuroectoderm and contributing to Primitive Streak fates. We find that Wnt signalling does not suppress neural development as previously thought and that it aids both fates in a context dependent manner. Our results suggest that as cells exit pluripotency they are endowed with a primary neuroectodermal fate and that the potency to become endomesodermal rises with time. We suggest that this situation translates into a “race for fates” at the level of single cells in which the neuroectodermal fate has an advantage.

## Introduction

Embryonic Stem (ES) cells are clonal populations derived from early mammalian blastocysts which have the ability to self renew as well as the capacity to differentiate into all embryonic cell types of an organism in culture and contribute to the normal development of an embryo into an organism, i.e. they are pluripotent [1,2,3]. Controlled application of defined cocktails of signalling molecules has been used as a means to differentiate ES cells into specific tissues, e.g. cardiac, blood, pancreatic cells, lung, gut, neurons [4,5,6,7,8,9] and, in some instances, into organ like structures resembling the in vivo counterparts [10]. However, despite notable successes, these protocols remain tinkering exercises performed with limited understanding of the routes and mechanisms that control differentiation. Furthermore, perhaps for this reason, questions remain as to the similarities and differences between the events in ES cells and in the embryo [11,12]. Therefore, gaining insights into the mechanisms of differentiation in culture will have an impact in our ability to harness the potential of these cells.

The experimentally controlled differentiation of pluripotent cells represents a good model system to understand the general process of fate decision making in development [4,13]. In the embryo, the blastocyst differentiates into the epiblast where, after implantation, within a single cell layered epithelium and under the influence of spatially organized signalling centres, cells are assigned to one of two fates: 1) an anteriorly located neural primordium or neuroectoderm (NECT), that will give rise to most of the brain, the anterior nervous system and epidermis, and 2) a group of cells in the proximal posterior region that will contribute to a dynamic structure, the Primitive Streak (PS), which acts as a seed for the endoderm and the mesoderm [14,15,16]. The spatiotemporal organization of the molecular components enabling and implementing the specification of these cell types are well known [17] and the sequence of events can be mimicked in culture [12,18], albeit in a spatially disorganized manner and often with low yields of cells of a particular fate. Notwithstanding these experiments, the details of the interactions between signalling and transcriptional networks mediating this decision remain poorly understood.

A widely held view of the differentiation of mammalian ES cells entertains that, as cells exit pluripotency, they choose between fates that resemble NECT and PS from a naïve state under the influence of their local signalling environment that enforces a rearrangement of the pluripotency network [19,20]. In this process, Retinoic Acid (RA) is deemed to promote NECT whereas Wnt/β-Catenin signalling might bias the decision towards the PS like fate [19]. However, NECT can emerge in the absence of RA [21,22] and Wnt/β-Catenin signalling has been reported to be required for the differentiation of neural precursors [23,24]. Therefore these observations raise questions on the prevalent model and led us to analyze in detail the influence of external signals on the early stages of differentiation of ES cells into either NECT-like or endomesoderm fated PS-like (from hereon referred to simply as NECT and PS).

Our observations support suggestions that ES cells have an intrinsic competence towards the NECT fate [22,25,26] and show that, after two days of undirected (neutral) differentiation, they enter a transient period of competence to become mesendoderm. Competence towards this fate emerges progressively and requires the downregulation of the pluripotency factor Nanog. We find no evidence to support the widespread notion that Wnt/β-Catenin signalling suppresses NECT [6,21,27,28,29,30] and instead find that, within the period of competence, it potentiates NECT and mesendoderm in a context-dependent manner. In the presence of high levels of Activin/Nodal and BMP, Wnt/β-Catenin signalling promotes mesendoderm whereas in their absence it produces anterior neurons.

## Results

### The exit from pluripotency establishes competence to differentiate

We have monitored the NECT/PS fate choice in differentiating ES cell populations using a Sox1::GFP reporter line for NECT, and a Brachyury::GFP (T::GFP) line for PS. The expression of both is negligible in self renewing conditions (Serum + LIF (SL), LIF + BMP (data not shown) or 2i + LIF) but can be detected when cells differentiate [31,32,33,34,35] (Fig. 1A). Low density growth in N2B27, which favours neural fates, elicits Sox1::GFP expression in about 70-90% of the cells, while growth in Activin and CHIR99021 (Chi, a GSK3 inhibitor that acts as an agonist of Wnt signalling; AC) activates T::GFP in about 30-50%, of cells (Fig. 1A).

**Figure 1:**
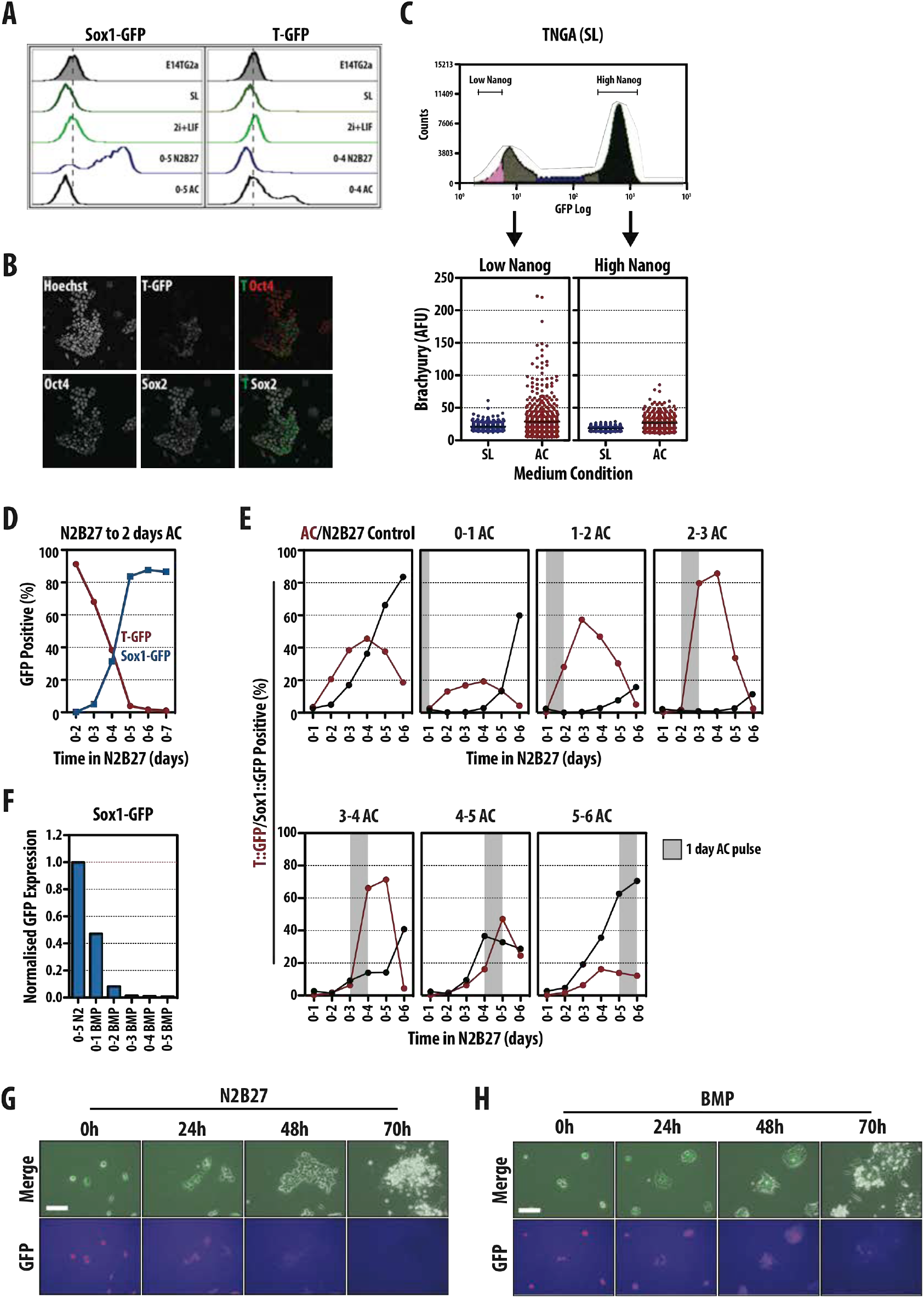
The exit from pluripotency determines the ability of mESCs to differentiate. (A) Sox1::GFP (left) or T::GFP (right) mESCs exposed to serum and LIF (SL), 2i and LIF (2i+L), N2B27 or Activin and Chi (AC) for the indicated durations and GFP expression analyzed by flow cytometry. Hashed vertical line bisecting the population profile plots indicates the peak maximum of the negative control (B) T::GFP mESCs differentiated in AC for 2 days (0-2), immunostained for Hoechst, Oct4 and Sox2 and imaged by confocal microscopy. Scale bar indicates 100 µm. (C) TNGA mESCs cultured in SL and FACS sorted into low- (indicated in pink) and high- (indicated in dark blue) expressing populations (top) were re-plated in AC conditions for 2 days, immunostained for Brachyury and analyzed by confocal microscopy (bottom). TNGA cells cultured in SL conditions served as a negative control for Brachyury immunostaining. (D) T::GFP (red) and Sox1::GFP (blue) mESCs were differentiated for 2 days in AC conditions after exposure to N2B27 for the indicated durations. GFP expression was measured by flow cytometry. (E) T::GFP (Red) or Sox1::GFP (black) mESCs plated and treated with N2B27 for 6 days with single, 1 day pulses of AC on days indicated above graphs. For the control, T::GFP or Sox1::GFP cells were incubated with 6 days of either AC or N2B27 respectively. Grey bar indicates period of AC pulsing. (F) Sox1::GFP mESCs treated with 1ng/ml BMP4 for durations indicated or 5 days (0-5) in N2B27. GFP expression measured by flow cytometry and fluorescence values displayed are normalized to the N2B27 control. (G,H) Live-cell imaging of TNGA cells in N2B27 alone (G) or supplemented with 1ng/ml BMP4 (H). Scale-bar represents 100 µm.

The relatively low yield of cells expressing T::GFP in PS differentiation conditions was surprising. The differentiated population contains a mixture of T::GFP-positive and negative cells interspersed with some that express Nanog (data not shown), Oct4 and Sox2 (Fig. 1B), indicating that some cells did not exit pluripotency and suggesting an explanation for the low percentage of T::GFP expressing cells. In self-renewing conditions ES cells exhibit a variegated differentiation potential with low Nanog expressing cells primed for differentiation [36,37]. Thus, it might be that only these cells can respond to AC. To test this we used TNGA cells, a Nanog::GFP reporter line that allows the identification of low Nanog expressing cells by their levels of GFP [36]. Sorting cells with high and low Nanog::GFP expression and exposing them to AC revealed that only the low Nanog population is capable to develop T::GFP expression effectively (Fig. 1C), i.e. lowering of Nanog might be a prerequisite for PS differentiation. In agreement with this, the percentage of T::GFP cells that respond to AC increases with the time of exposure to N2B27, reaching a peak of more than 80% after two days when all cells in the population have low levels of Nanog (Fig. 1D,E). The rise in the percentage of T::GFP positive cells is, in addition, highly correlated with a decrease of Sox1::GFP positive cells (correlation coefficient of −0.97, Fig. 1D). Treating cells to a short pulse of AC (1 day) after increasing durations of N2B27 exposure revealed that cells are most sensitive to PS-inducing stimuli after two days (Fig. 1E); as this duration was extended, cells began to favour a NECT fate (Fig. 1E).

To test if there was a similar change in the competence for NECT over the first two days of induced differentiation, we made use of the ability of BMP signalling to inhibit this event [38,39,40,41,42]. Treatment of ES cells with 1ng/ml BMP4 for different periods of time (Fig. 1F) confirmed previous reports: treatment during the first day of differentiation vastly reduced Sox1::GFP expression, whilst exposure for longer periods, and even for the second day of differentiation only, eliminated Sox1::GFP expression (Fig. 1F). However this effect might not be related to an inhibition of the NECT fate, but rather to an enhancement of pluripotency [38,43]. Consistent with this we observe that when TNGA cells are exposed to 1ng/ml BMP4 and filmed over time, they retain Nanog::GFP expression for about two days before abruptly differentiating (Fig. 1G,H); this is confirmed in large population studies using flow cytometry (Fig. 1F and see below).

Altogether these results suggest that at the exit from pluripotency, ES cells respond differently to NECT and PS differentiation signals. Our results agree with previous observations that commitment to NECT, as reflected in Sox1::GFP expression, does not appear to require specific inputs other than those produced by the cells [21,22,44] and may be latent in self renewal conditions [25,26]. On the other hand we notice that the ability to respond to AC emerges as cells shut down the pluripotency network. In both cases, commitment to a particular fate requires a downregulation of Nanog and we observe that only after 2 days of undirected differentiation is the population, as a whole, competent to respond to AC (Fig. 1D,E,G).

### A balance of Wnt, Nodal/Activin and BMP signalling controls fate decisions at the population level during the exit from pluripotency

There is evidence that, in addition to BMP, Activin/Nodal and Wnt/β-Catenin signalling can suppress NECT specification during ES cell differentiation [6,21,27,28,29,30]. In agreement with this, treatment of mES cells induced to differentiate in N2B27 for 5 days with Activin (Act), BMP4, or the GSK3 inhibitor CHIR99021 (Chi) to induce Wnt/β-Catenin signalling, results in inhibition of Sox1::GFP expression (Fig. 2A). Whereas, inhibition Activin activity with SB431542 (SB43), BMP activity with DMH1 or β-Catenin activity with the Tankyrase inhibitor XAV939 resulted in enhanced Sox1::GFP expression (Fig. 2A). To test the impact of each of these signals and their interactions on the NECT/PS decision during the exit from pluripotency, we treated ES cells with agonists and antagonists of each pathway for the first two days of differentiation, then transferred them to either N2B27 (3 days) or AC (2 days) and recorded Sox1::GFP or T::GFP expression (Fig. 2).

**Figure 2:**
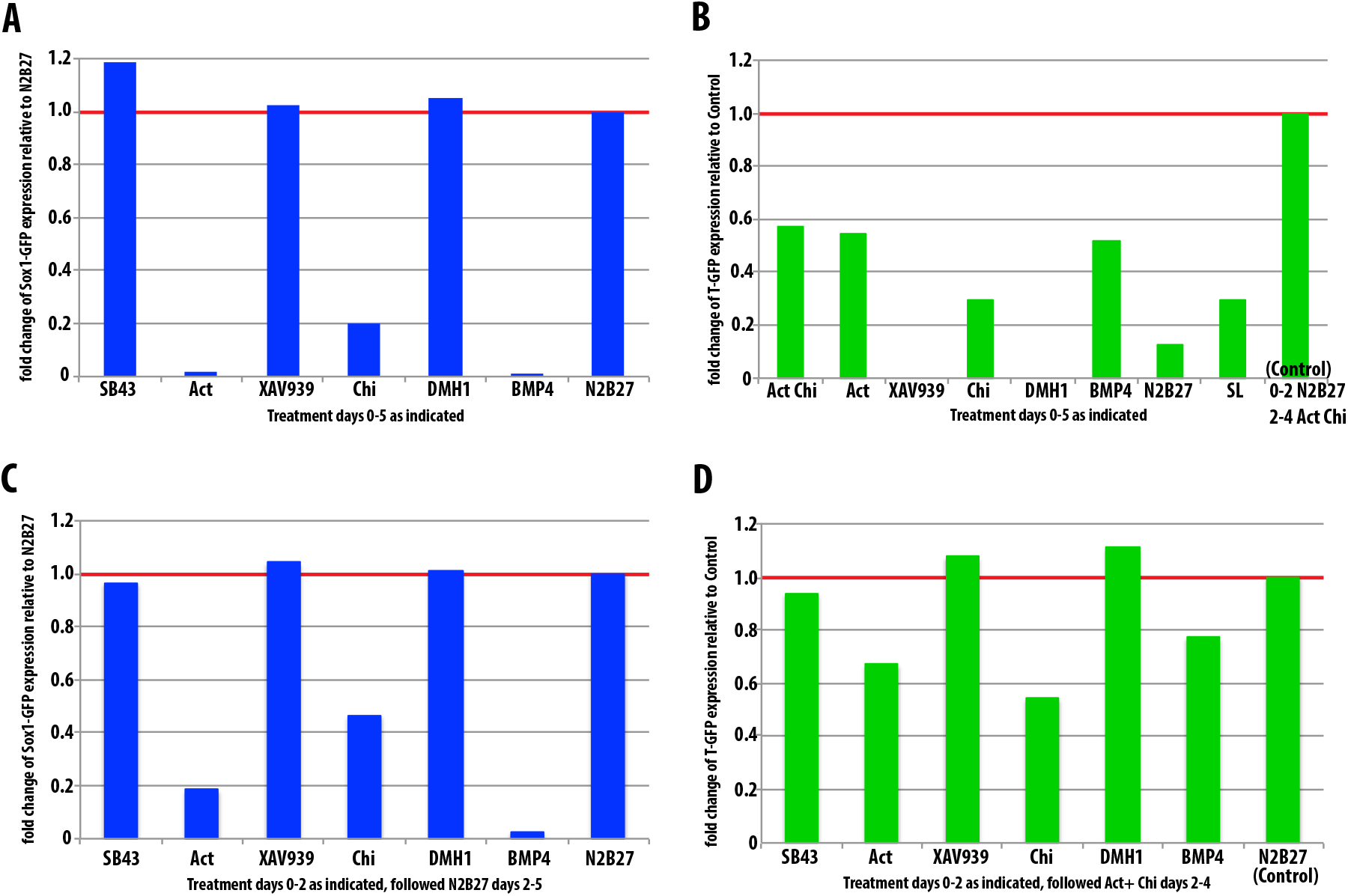
Effects of signalling during the exit from pluripotency. (A, B) Proportion of GFP positive cells in Sox1::GFP (A) or T::GFP (B) cells exposed to the indicated factors for 5 days. C, D) Proportion of GFP positive cells in Sox1::GFP (C) or T::GFP (D) cells exposed to the indicated factors for 2 days prior to switching media to N2B27 for 3 days (A, Sox1::GFP) or AC conditions for 2 days (B, T::GFP). Expression of GFP was measured by flow cytometry. All data presented here is normalized to 5 days in N2B27 (Sox1::GFP) or 2 days N2B27 followed by 2 days AC conditions (T::GFP).

Activation of Wnt/β-Catenin signalling with Chi, either for the first two days or for the period of differentiation, resulted in a suppression of Sox1::GFP expression (Fig. 2A, C), whereas inhibition of β-Catenin activity with XAV939, results in some apoptosis, but the remaining cells exhibit a robust activation of Sox1::GFP (Fig. 2A,C). The effects of Wnt/β-Catenin signalling on T::GFP expression depend on the period and timing of exposure. Even though β-Catenin is necessary for T::GFP expression, treatment with Chi for the first two days before the exposure to PS-promoting signals (AC), reduces, rather than enhances the number of T::GFP expressing cells (Fig. 2D). Furthermore, exposure to XAV939 during the exit from pluripotency, enhances the response to AC. The effects of Activin signalling followed a similar pattern: exposure to Act during the first two days of differentiation inhibited Sox1::GFP expression (Fig. 2C) and to lesser extent, T::GFP expression (Fig. 2D). On the other hand, inhibition of Activin signalling during this period with SB43, led to an increase in Sox1::GFP and to a slight but detectable reduction of T::GFP expression by exposure to AC (Fig. 2D). In all cases the effects were less pronounced the shorter the exposure of the cells to the signal modulators (Fig. 2 Supplementary). Tests of the interplay between the two pathways on Sox1::GFP expression (Fig. 2 Supplementary) indicate a predominant role of an inhibitory function of these signals, with the effects of Activin and Chi being dominant over the loss of function of either pathway. We noticed that, in contrast to the effects of BMP, one day exposure to either Activin or Chi was not sufficient to suppress Sox1::GFP expression (Fig. 2 Supplementary).

Altogether these results suggest that, in agreement with published reports, both Wnt/β-Catenin and Activin/Nodal signalling can suppress NECT fate during the first two days of differentiation. However, as β-Catenin signalling has been shown to promote pluripotency [45,46,47,48,49,50], there is a possibility that, as in the case of BMP, its effects on Sox1::GFP expression reflect a function in the maintenance of pluripotency rather than on neural differentiation. Consistent with this flow cytometric analysis and live imaging of ES cells (Figs. 2 and 3A-D) show that the effects of Wnt/β-Catenin signalling on the early stages of differentiation are a consequence of its effects on the stability of pluripotency: activation of β-Catenin maintains Nanog::GFP (TNGA cells) and REX1::GFP reporters, whilst inhibition with XAV939 promotes their downregulation (Figs. 2 and 3A-D).

**Figure 3:**
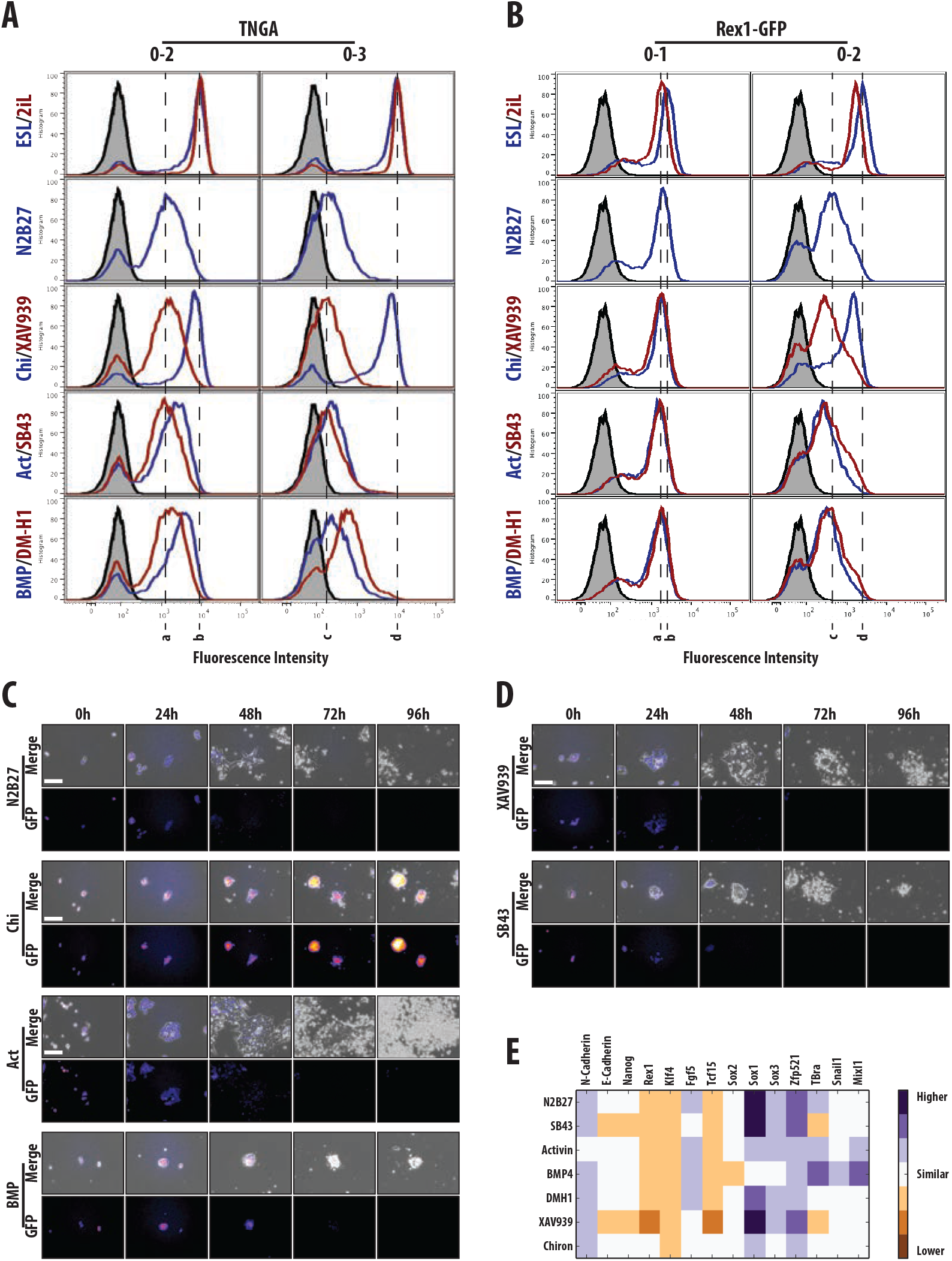
A fate restriction point at the exit from pluripotency. (A,B) Population profiles of cells analyzed by flow cytometry from either (A) Nanog::GFP (TNGA) or (B) Rex1::GFP mESCs after 2 (0-2) or 3 (0-3) days in the medium indicated (SL, 2iL, N2B27, Chi, XAV939, Activin (Act), SB43, BMP, DM-H1). Hashed vertical lines correspond to the peak maximum of the fluorescence from cells in either N2B27 (a,c) or SL (b,d) conditions at each time-point. (C) Live-cell imaging of TNGA mESCs differentiated for 96 h in N2B27, Chi, Act or BMP. (D) Live-cell imaging of TNGA mESCs differentiated for 96 h in either SB43 or XAV939. Scale-bar in all live-cell imaging experiments indicates 100 µm. (E) Sox1::GFP cells were grown in the indicated medium for 2 days and a further day in N2B27 prior to RNA extraction and RT-qPCR analysis for the indicated genes. Data normalized to the house-keeping gene Ppia (see supplemental data for Fig. 3) and displayed as values higher, similar or lower than the SL control.

The notion that the impact of β-Catenin on the early stages of differentiation reflects its activity in promoting pluripotency is confirmed by its effects on PS differentiation. Although Wnt/β-Catenin signalling is required for PS fate, treatment of ES cells with Chi before exposing them to AC decreases rather than increases, the proportion of cells that develop as PS (Fig. 2D). This surprising observation can only be understood in terms of the effect that β-Catenin has in promoting pluripotency which will thus reduces the possibility to commit to any differentiation route.

Whereas exposure to Activin or SB43 during the exit from pluripotency does not alter the dynamics of the Nanog::GFP or Rex1::GFP reporters with respect to the N2B27 control (Fig. 3A–D), exposure to Activin suppresses Sox1::GFP and, to a lesser degree, T::GFP expression (Fig. 2C–D). The effect on NECT cannot be explained in terms of effects on pluripotency since Activin/Nodal promotes exit from pluripotency ([51] and Fig. 3) and therefore must reflect an active suppression of the NECT fate; the effects of SB43 on Sox1::GFP confirm this possibility (Fig. 2A,C). On the other hand, the reduction of T::GFP expression upon exposure to Activin (Fig. 2D) is surprising and might reflect a delayed differentiation associated with the requirement for Activin/Nodal in the maintenance of the EpiSC state [51,52] i.e. the exposure to Act induces a slow transition through an Epiblast like state and a delay in cell fate decisions through a pause in an Epi-like state.

### The impact of signals on gene expression

To better characterize the impact of the signalling pathways on the exit from pluripotency, we monitored the expression of genes associated with pluripotency (E-Cadherin, Nanog, Rex1, Klf4), the exit from pluripotency (Tcf15), the epiblast (Fgf5), neural (N-Cadherin, Sox1, Sox2, Sox3 and Zfp521) or PS differentiation (T/Bra, Snail, Mixl1) under different conditions. Cells were kept for 2 days in the different conditions and cultured for a further day in N2B27 before their state being assessed (Fig. 3E). The results provide some insights into the action of the different signals. For example they confirm that the effects of β-Catenin on the commitment to different fates are due to its effects on pluripotency: Chi, which increases ß-Catenin, maintains the levels of all pluripotency markers with little effect on differentiation, whereas XAV939, which reduces ß-Catenin, promotes differentiation, with a bias towards NECT (Fig. 3E) in the absence of additional signals. The effect of BMP is more subtle and gene expression confirms that it delays, rather than suppresses, differentiation; in this experiment cells seem to be in an Epi-like state but the high levels of expression of T/Bra, Snail and Mixl1 in these cells suggest that they have a strong bias towards PS (see also [43]). The profile also confirms the effects of Nodal/Activin on NECT differentiation but also provides some support for their effect in delaying differentiation through a maintenance of an Epi-like state as reflected in the high levels of Fgf5 and the mixture of neural and PS gene expression within the population.

Altogether these observations show that, as cells exit pluripotency, they integrate several signals in a time dependent manner. They also emphasize the need to take into account the dynamics of the pluripotency network in cell populations when considering the impact of different signals in the process of differentiation. Cells will respond to individual signals depending on the state they are in. In an ES culture maintained in Serum and LIF, there is a mixture of cells with varied differentiation potential. Cells with high levels of Nanog are pluripotent and both BMP and β-Catenin signalling will enhance this state, while Activin/Nodal and FGF/ERK signalling will contribute to loss of Nanog expression and thereby promote differentiation [51,53,54,55,56]. On the other hand, the low Nanog expressing population is an heterogeneous mixture of cells in different states: some primed to return to the high Nanog state, some in an epiblast (or Epi Stem Cell) like state and some differentiating or even already differentiated. This population with low levels of Nanog exhibits higher levels of Wnt/β-Catenin transcriptional activity [45,57] and its response to signals will depend on the particular state of the cell. For those primed to differentiate, the combined activity of Wnt, Activin, BMP and FGF will favour differentiation and, specifically, differentiation into PS fates. Our experiments also support the notion that NECT might not need specific external signals whose effect is, for the most part, to induce a PS fate (Fig. 2, Fig. 2 supplementary data, Fig 3E).

### A fate restriction/commitment point at the exit from pluripotency

Our results support the contention that BMP delays differentiation [38,43] and show that this effect is dominant over pro-differentiation signals as BMP suppresses the effects of SB43 and XAV939 on the exit from pluripotency (Fig. 2 supplementary data). However, BMP does not abolish differentiation. Monitoring persistent exposure of ES cells to BMP reveals an abrupt loss of Nanog expression at day 3 (Fig. 3A,C), and the emergence of T::GFP expression (Fig. 2B). Furthermore, BMP boosts the effects of Activin and Chi on the expression of T::GFP, in particular in the context of differentiation into PS derivatives [9,35,58,59]. This effect is more obvious when BMP is applied after two days of differentiation and results in a total suppression of Sox1::GFP expression without reverting the cells to a pluripotent state (Fig. 5 and Fig 5 supplementary data). Taken together these observations highlight a change in the responsiveness of the differentiating population of ES cells after 2/3 days of differentiation in culture. The most significant change is the emergence of a competence to respond to PS-promoting signals, which emerges at this time and lasts for two days (Fig. 1D,E).

A number of studies show that at day 3 of differentiation, ES cells go through a state similar to that of Epiblast stem cells, which can be mapped onto the post-implantation epiblast E5.0-E6.0 [38]. It has been further suggested that at this moment cells are primed for both NECT and PS and that they choose between the two fates [19,60,61]. This is supported by our observation that at this point cells lose the ability to return to the naïve state (not shown) and can be seen most clearly with live-cell imaging and the FACS profiles where we observe morphological and dynamical changes in the cells around this time (Fig. 3A–D). [We need to discuss these results; at least to mention them].

### Gene expression during exit from pluripotency

To understand the molecular basis of fate choice during mouse ES cell differentiation, we monitored the expression of a collection of genes associated with pluripotency and specific lineages as well as elements of various signalling pathways in ES cells as they exit pluripotency. In a first experiment, cells in LIF and BMP were allowed to differentiate in N2B27 for five days and gene expression was monitored both in the population (Fig. 4A,B,C) and at the level of single cells (Fig. 4D,E). Initial and strong changes in expression are largely restricted to pluripotency-related genes but lineage affiliated gene expression emerges as cells begin to differentiate (Fig. 4A); in addition, we can see that at days 2 and 3, the number of genes expressed is maximal (data not shown). A survey of specific genes confirms that the exit from pluripotency is associated with a downregulation of genes associated with the pluripotency network (Nanog, Oct4, Sox2, Klf4, Esrrb) and that at about day 3 the cells go through an Epi-like state as characterized by the expression of Fgf5 and a transient rise in Nanog expression (Fig. 4A). In terms of differentiation, whilst genes associated with NECT (e.g. Sox1, Sox2, Sox3, Pax6, Pax7, Otx2) are expressed as the cells begin to differentiate and continue to be expressed during the differentiation period, genes associated with PS (e.g. Foxa2, Gsc, T/Bra, Eomes, Mixl1, Snail, Tbx6, Wnt3) emerge around day 3 (Fig. 4A). This study also confirms the rise of Wnt/β-Catenin transcriptional activity described before [45] in the expression of direct targets, e.g. Dkk1 and Axin2, which rise during day 3 (Fig. 4A).

**Figure 4:**
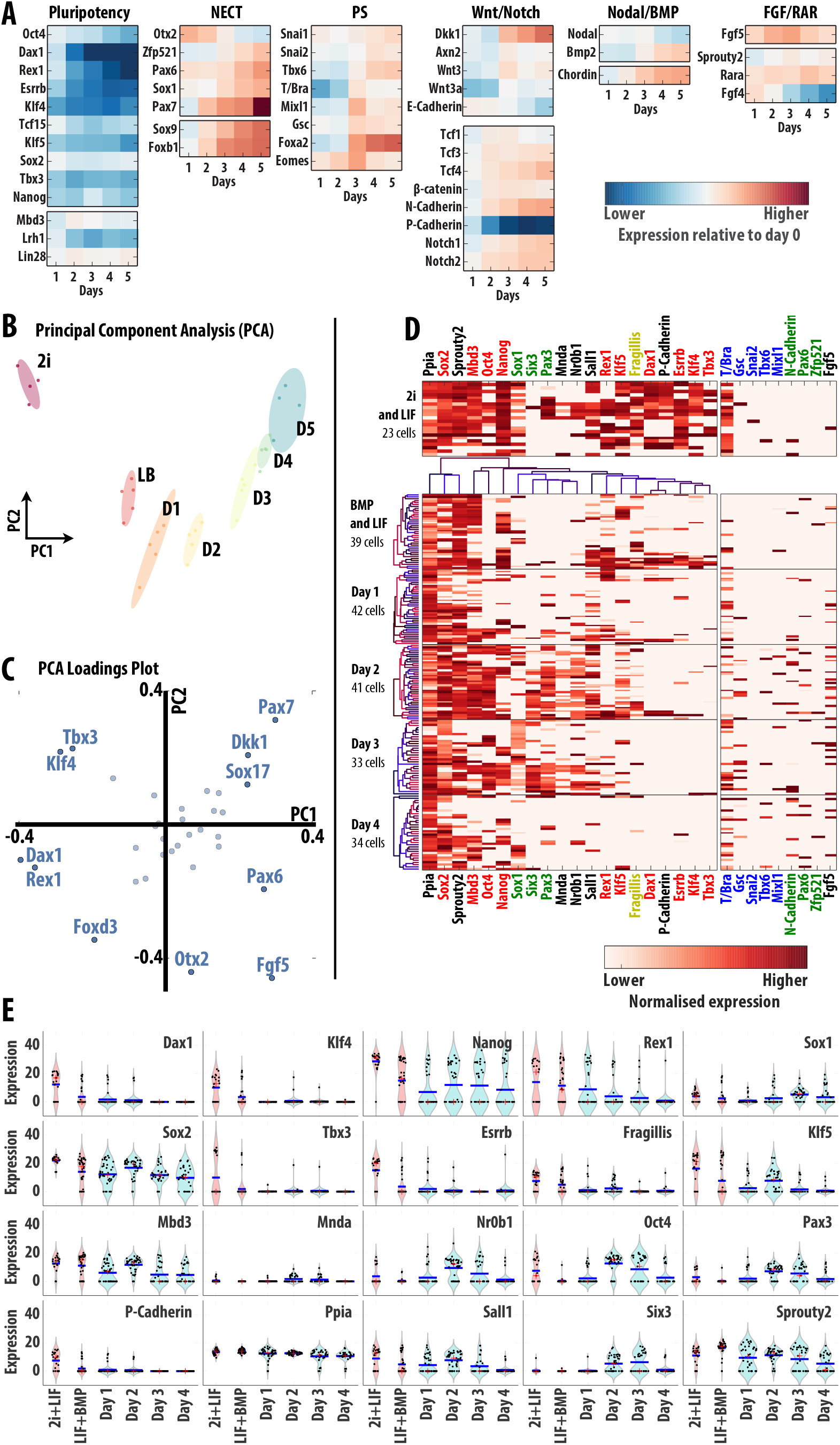
Dynamics of gene expression during the exit from pluripotency. Mouse ES cells cultured in LIF and BMP were transferred to N2B27 and allowed to differentiate for 5 days. (A) Average (bulk) expression levels of several genes associated with pluripotency, NECT and PS fate restriction as well as targets for Wnt/Notch, Nodal/BMP and FGF/RAR signalling were measured daily using fluidigm quantitative RT-PCR. Color-coded values indicate the log-fold change values with respect to the initial LIF + BMP condition expression levels measured by RT-PCR and normalized to a housekeeping gene. (B) Principal Component Analysis of the bulk expression profiles of 34 genes for 2i, LIF and BMP and the 5 days of differentiation in N2B27 normalized to Gapdh. Each single point indicates a technical repeat. (C) Loadings plot of the PCA indicating the genes that contribute the most to the first two components. (D-E) Expression levels of over 90 genes analyzed using fluidigm techology and single-cell RT-PCR over the 5-day differentiation experiment. (D) Heat maps of single-cell expression levels normalized to Gapdh (each row represents an individual cell) for the most significant genes. Genes (in columns) were clustered according to the pairwise similarities in their expression levels. Additional genes of interest are also displayed in the right-most panels. Data for 2i and LIF conditions is also shown on the panel on top. (E) Violin plots of the data in D showing kernel density estimates of the mRNA distribution (clear blue) together with single cell expression levels (black dots), the population average (blue line) and median (red dot).

Principal Component Analysis (PCA) of the expression profiles of 34 genes during the exit from pluripotency in undirected differentiation conditions (Fig. 4B,C and Fig. 4 Supplementary Data A-F) provides additional insights into the process. The first component (component of maximum variance, PC1 in Fig. 4B) clearly delineates a differentiation axis, with all differentiation stages clustered in chronological order from left to right. This first component accounts for more than 60% of the total variance (Fig. 4 Supplementary Data B) and is mainly defined by genes of the pluripotency network, such as Rex1, Dax1, Klf4 and Tbx3 (Fig. 4C and Fig. 4 Supplementary Data C), with a contribution of others like Fgf5 associated with the transient Epi-like state. The second component, PC2, which can be interpreted as the main axis of change in gene expression once the effect of loss of pluripotency genes (PC1) has been removed, amounts to about 15% of the total variance. Interestingly, this component highlights an inflection point on day 3 of differentiation (Fig. 4B; this discontinuity is even more evident when 2i conditions are not considered for the analysis, see Fig. 4 Supplementary Data F). Genes strongly contributing PC2 include some that are highly expressed during the first two days: Otx2 and Foxd3; as well as genes that are very lowly expressed in 2i, all of which appear to be lineage affiliated (Fig. 4C and Fig. 4 Supplementary Data C).

The PCA analysis highlights that the loss of pluripotency gene expression is a major driver of the differentiation process. This is confirmed by the analysis of gene expression in single cells with the core elements of the pluripotency network (Essrb, Oct4, Dax1, Klf4, Sox2, Nanog, and Rex1) which are amidst the top 15 genes driving the change (Fig. 4D,E, and Fig. 4 Supplementary Data G). An interesting observation in this list is Mbd3, a core member of the NurD complex, whose expression decreases during development and which our analysis suggests it is a driver of the process. Another interesting one is P-Cadherin, which has been highlighted before [45]. As differentiation progresses, single cells start to favour the high expression of genes associated with the NECT (Fig 4D and Fig. 4 Supplementary Data G).

### Signals and a decision point for commitment and differentiation

After three days in N2B27, cells are unable to return to the naïve state and we presume that they will differentiate according to their local signalling environment. We have explored this by exposing cultures to different signals and signal combinations from day 2 of differentiation (Fig. 5). As expected, exposure to Activin or BMP promotes T::GFP expression and a PS fate (Fig. 5A and Fig. 5 Supplementary data) while suppression of Activin signalling (with SB43) or BMP (with DMH1) promotes Sox1::GFP expression and the NECT fate (Fig. 5B). Surprisingly, we find that exposure to Chi from this time promotes both Sox1::GFP and T::GFP expression in a context dependent manner: it both enhances the effects of Activin on the PS fate and the effect of SB43 on the NECT (Fig. 5C).

**Figure 5:**
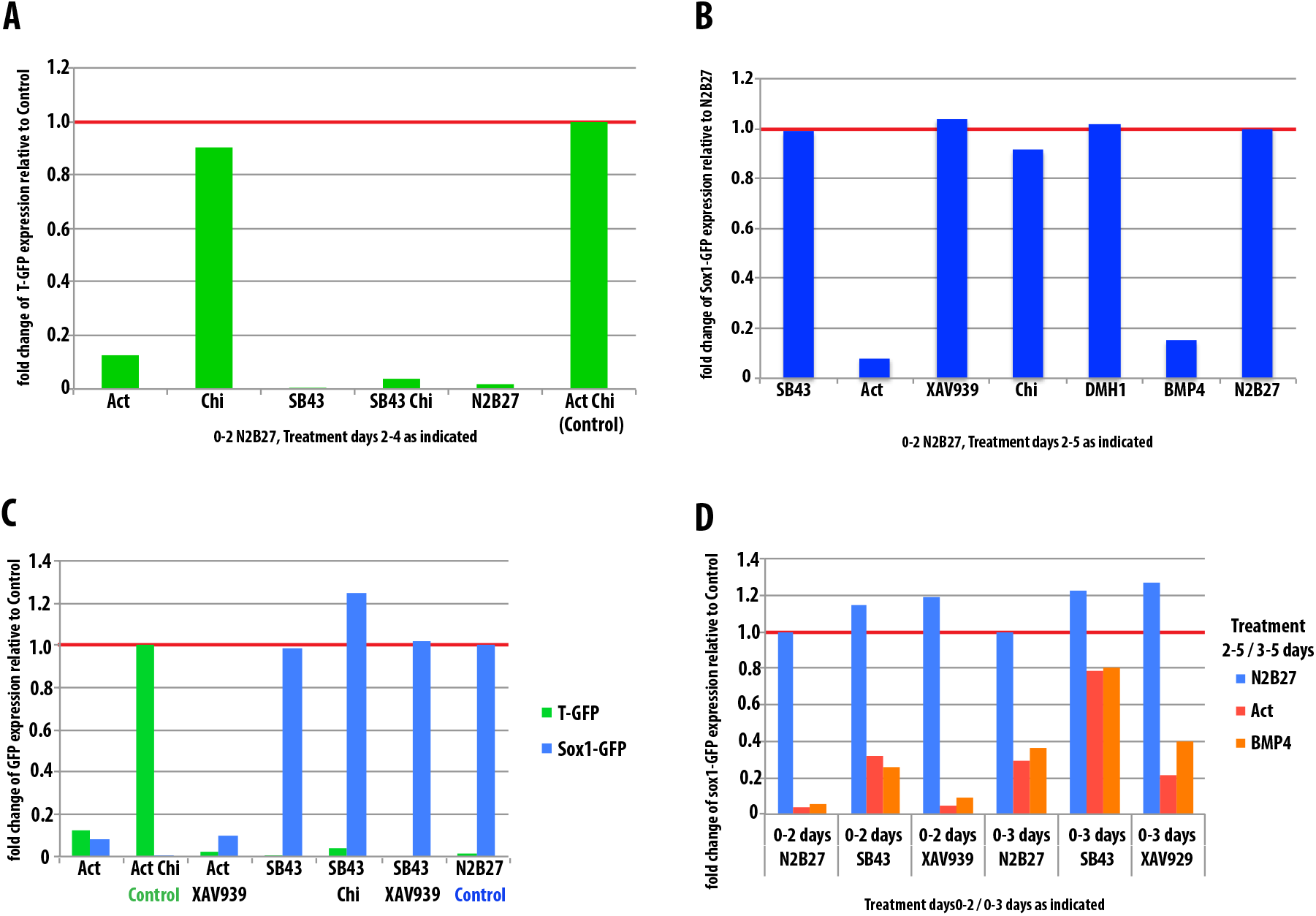
Activin activity but not Wnt/β-Catenin activity controls fate decisions between PS and NECT. (A-C) T::GFP (A) and Sox1::GFP (B, C) cells were grown in N2B27 for 2 days prior to transferring to the indicated conditions for 2 days (T::GFP) or 3 days(Sox1-GFP). GFP expression was assessed by flow cytometry and data normalized to control conditions (0-2 days N2B27, 2-4 days AC for T::GFP, 0-5 days N2B27 for Sox1::GFP) and presented in bar charts. (D) Sox1::GFP cells were plated in either N2B27, SB43 or XAV939 for the first 2 or 3 days of culture and then switched to N2B27, ACT or BMP4 for the remaining period. GFP expression was assessed by flow cytometry on day 5. Data is displayed as a bar chart and has been normalized to control conditions (0-5 N2B27).

Altogether these results indicate that the competence of cells to respond to AC and acquire a PS fate is maximal after 2 days of undirected differentiation (Fig. 1E), whereas the competence for NECT is present from the initial stages of differentiation but declines probably due to exposure of the cells to paracrine Activin/Nodal signalling. Inhibition of both Activin with SB43 and Wnt/β-Catenin signalling with XAV939 enhance both the proportion and rate of cells exiting the pluripotent state (Fig. 3A,B,E) and also leads to an enhanced population of Sox1::GFP. This increase could be attributed to an enhanced rate of exit from pluripotency, which would confer an advantage to the NECT fate, or to an enhanced commitment to NECT fate. To test this, cells were exposed to SB43 or XAV939 for the initial 2 or 3 days of differentiation before switching to either N2B27, Activin or BMP for the remainder of the 5 day assay (Fig. 5D). Whereas populations of cells initially exposed to XAV939 resembled those cultured in N2B27 (with respect to the proportion of Sox1::GFP-positive cells), cells initially grown in SB43 showed an enhanced commitment to the NECT fate and the longer the exposure the greater the enhancement (Fig. 5D). This result emphasizes that while Nodal/Activin signalling suppresses NECT with little effect on pluripotency, the apparent suppression of NECT fate mediated by Wnt/ß-catenin signalling is an indirect effect of its activity in promoting pluripotency.

In summary, whereas Wnt/β-Catenin signalling regulates the dynamics of the exit from pluripotency, it has little effect on the fate of the cells during the first two days of differentiation. On the other hand, Nodal/Activin controls both the rate of differentiation and the fate of the cells. At day 3 of differentiation in culture, cells have a competency to become committed to either of the fates under the influence of the signalling environment. Activin and BMP signalling promote PS and inhibit NECT whereas Wnt/β-Catenin signalling appears to play a role of enhancement of both fates in the context of the decision.

### Discussion

We have analyzed the role of signalling in fate assignment during the early stages of differentiation of mouse ES cell populations in culture. While there are many studies of ES cell differentiation into particular cell types, there is no integrated analysis of the kind that we have performed here. Our observations reveal the importance of the state of the cell when responding to signals, particularly to those that promote both self-renewal and differentiation like Wnt and BMP. Both signals have been suggested to suppress neural differentiation [6,21,27,29,38,40,41,42,62], however we find that at the exit from pluripotency, their effects on differentiation are mediated, mostly, through their effects on pluripotency. Our results support the conclusions that BMP delays the exit from pluripotency [38,43] whilst simultaneously priming the PS fate [34,35,43]. In the case of Wnt/β-Catenin, we find that the observed suppression of neural fates is an indirect consequence of its effects on pluripotency [45,63]. High levels of β-Catenin maintain cells pluripotent but this activity is not stable. Over time, cells exit pluripotency and when they do, in the presence of high levels of β-Catenin they adopt the PS fate. We find that in cells committed to differentiation on which BMP suppresses the neural fate, Wnt/β-Catenin signalling promotes both NECT and PS fates in a signalling context dependent manner. While its effect on PS is in agreement with well-established observations [9,12,64], the effect on NECT development is, at first sight, surprising. However there is evidence that Wnt/β-Catenin signalling promotes the differentiation of neural precursors [23,24] and some of our observations are likely to reflect this activity.

Our results suggest that the effects of β-Catenin depend on the signalling context it is embedded in. For example, in the presence of high levels of β-Catenin, it is the levels of Activin that determine the fate of the cells: high levels, as provided experimentally, promote endoderm, whereas lower levels promote the development of mesoderm. It is also possible that even within the anterior neural fate, which develops in the absence of Activin/Nodal signalling, the levels of β-Catenin might determine fates. In this context, it is of interest that an effect of β-Catenin (Chi) on NECT differentiation can be observed in Fig. 7G in [19] but is ignored by the authors. The lack of lineage specificity in the effects of Wnt/β-Catenin signalling contrasts with those of Nodal/Activin and BMP which promote PS and suppress NECT and are consistent with a proposed role of β-Catenin affecting the probability with which a cell adopts a fate rather than the implementation of such fate [65,66].

Our results support the notion that after two days of differentiation, ES cells enter a state in which they commit to particular fates [19]. This state is closely related to that of cells in the E5.5 epiblast and is reflected in the expression of Fgf5, the lowering of E-Cadherin, a transient rise in Nanog and the co-expression of genes associated with PS and NECT [38,57]. A prevalent view of this state of commitment is that cells make choices from a naïve state [19]; however, our results are consistent with a different view in which ES cells have a primary NECT fate which can be overridden for a short period of time by pro-PS signals. The notion of a “primary” or “default” NECT fate in ES cells has been suggested before [25,26] and is supported by the observation that in the absence any external signals ES cells will adopt a neural fate [21,22,44], as well as by our observation of the prevalence of NECT gene expression in self renewal and, particularly, during the early stages of differentiation. On the other hand, the competence to become PS emerges over time with a peak at days 3 and 4, and is associated with a decrease in Nanog expression, the loss of Oct4 expression and the emergence of β-Catenin transcriptional activity. These events can be observed in neutral differentiation conditions e.g. N2B27, where in the course of the time of the experiment, the levels of Activin/Nodal, BMP and Wnt can be seen to rise as reflected in the expression of PS genes e.g T/Bra, Mixl1, Eomes [34,67] (and here). These signals are likely to be diluted in the culture but can be effective in small patches and we can observe multiple fates arising over four or five days differentiation in N2B27 [57] (and here). However, the large dilution effects of the culture are likely to prevent significant long term differentiation of PS derivatives in these conditions and long term culture in N2B27 selects against PS cells. Conversely, addition of agonists of Activin/Nodal, BMP and Wnt selects for PS. Our results show that there is a window of opportunity to become PS and that if the cells do not adopt the PS fate in this period, they will become NECT.

Taken together our results lead us to surmise that the fate choice that cells face in culture is of the kind “IF NOT X then Y” i.e. “IF NOT PS then NECT”. Many of the pro- NECT effects of suppressing Wnt/β-Catenin or BMP signalling could be interpreted in this light: an early exit from pluripotency might give an advantage to commit to NECT before a build up of PS promoting signals. Consistent with this, filming of the exit from pluripotency and fate assignment shows that the cells that lower Nanog early have a higher probability to become neural (unpublished).

### Fate choices at the exit from pluripotency: a framework

In the context of published work, our results lead us to suggest a sequence of events and causal connections for the manner in which signals guide fate decisions at the exit from pluripotency in culture (Fig. 6A). To do this we integrate our results with a number of published facts:

1. A pluripotent population in standard self renewing conditions (Serum and LIF, BMP and LIF) is a dynamic mixture of three subpopulations: cells in the ground state, Epi stem cells and cells in transition between these two states; it also includes some differentiated cells [36,37,68].
2. The state of a cell in an ES cell population is determined by the levels of Nanog and β-Catenin which, in turn, determine the levels of free Oct4. Differentiation is determined by the Oct4:Nanog ratio: a high ratio it will promote differentiation [63,69,70,71].
3. The elements of the pluripotency network prime particular fates, with Oct4 priming all differentiation but then, together with low levels of Nanog, promoting PS and Sox2 promoting NECT [19,20].
4. Differentiation is associated with Wnt/β-Catenin signalling [45,57].
5. In N2B27 differentiation is driven by local signalling interactions between cells which can be overridden by external signals in experimental conditions.

**Figure 6:**
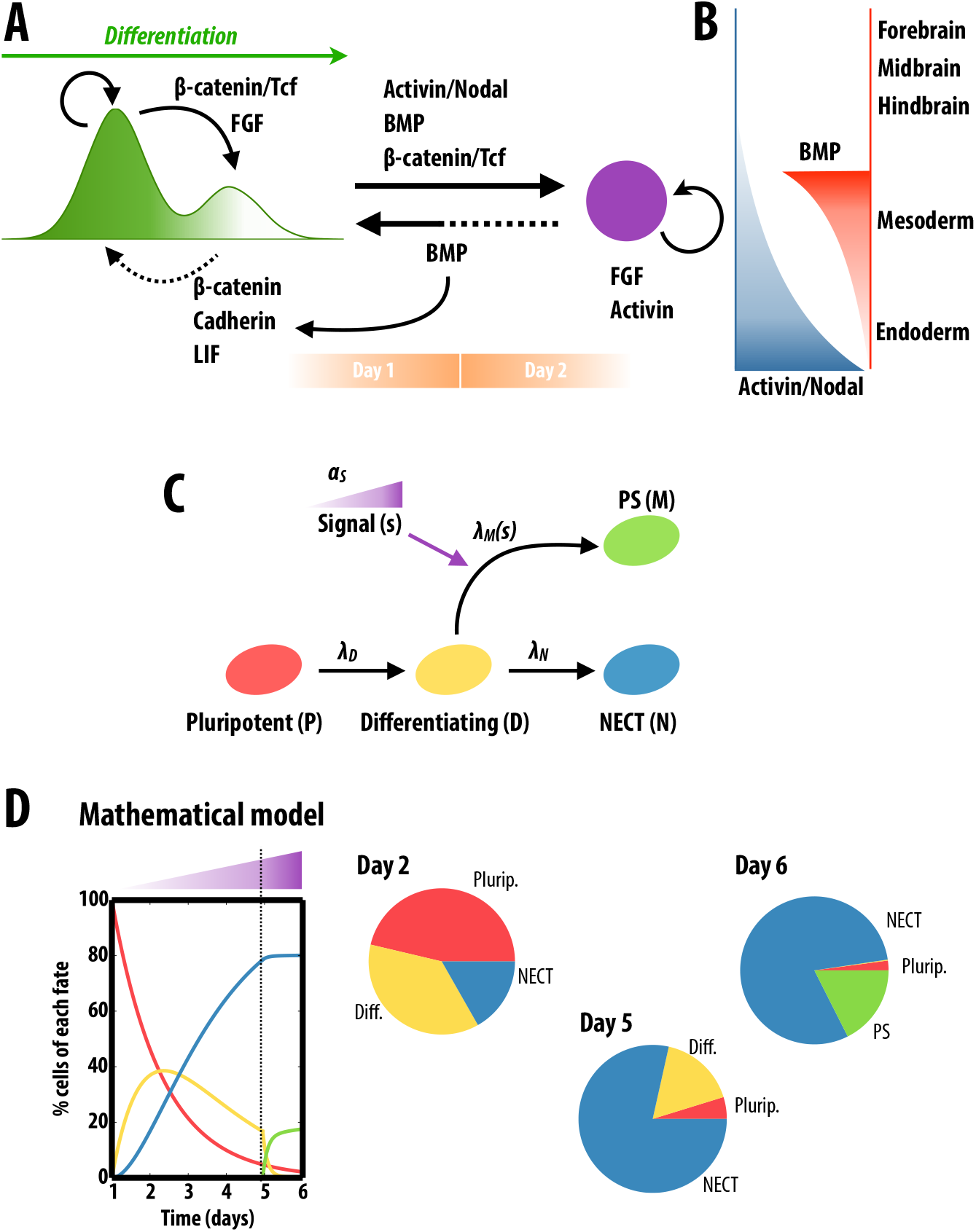
Fate choices at the exit from pluripotency. (A-B) Summary of interactions between signalling pathways and cell states during the exit from pluripotency and the commitment to different cell fates (for details see main text). Embryonic Stem cells self renew in a metastable state governed by the ratio of Nanog and Oct4 (indicated in the levels of green) and exhibit a bimodal distribution of the kind shown. The ratios are implemented by a combination of β-Catenin/E-Cadherin and inhibition of ERK signalling. When differentiation conditions are implemented in the culture ERK and Wnt/β-Catenin/Tcf signalling promote differentiation with levels of Nodal/Activin and BMP determining the fates of the cells. After two days of differentiation, cells enter in an Epiblast like state–maintained by FGF and Activin–where their fate is determined by the relative levels of BMP and Nodal/Activin as indicated. Wnt/β-Catenin signalling is required for all fates. (C) A simple model of cell-fate adoption describing the transitions from different cellular states: from pluripotent (*P*) to differentiating (*D*), and from the latter to either NECT (*N*) or PS (*M*); and the signalling effects in the transition rates accounts for the observed dynamics (D). Time traces of each cell fate as simulated by the model. In the initial stages of differentiation cells are more prone to adopt the NECT (blue) fate than the PS fate (green). As cells differentiate and the global levels of signal (e.g., Nodal/Activin) raise, the PS fate becomes dominant within the remaining differentiating cells. See Materials and Methods and Fig. 6 Supplementary Data for details of the mathematical model.

In self renewal conditions, a metastable balance of the activity of signal transduction networks maintains a ratio of Oct4:Nanog that fosters the activity of the pluripotency network and suppresses a high frequency of differentiation [37,63,72]. In this situation, BMP and Wnt/β-Catenin signalling in the context of low FGF/ERK signalling and high levels of E-Cadherin, play a pivotal role in this balance [43,45,46,73,74] and maintain the pluripotency network in a homeostatic steadiness. Triggering differentiation artificially and neutrally e.g. by placing the cells in N2B27, leads to a shift in the balance of signals and the Oct4:Nanog ratio leading to an activation of FGF:ERK and β-Catenin transcriptional activities [45,54,55]. This launches differentiation in the context of an intrinsic/primary NECT programme which firms up as the pluripotency network disassembles. During this period expression of BMP, Nodal and Wnt rises and we surmise that in N2B27 the levels of these signals increase in the medium and condition cells to the PS fate. However, it is clear that these signals are not sufficient to promote PS differentiation and that cells only respond effectively to them when they have disassembled the pluripotency network. At the moment we do not understand what this means in molecular terms i.e. what the molecular events are that link the disassembly of the pluripotency network and the commitment to particular fate, but our observations point out this correlation.

One can view the process as a race of two mutually excluding fates with two different triggering thresholds. The PS programme can suppress the NECT programme but not the other way around. However the suppression can only take place while the cells are deciding and not passed their commitment time. Furthermore, the competence to PS is not built into the system initially but arises during differentiation. With this in mind we have created a simple mathematical model that accounts for our observations (Fig. 6C–D, Fig. 6 Supplementary Data and see Materials and Methods). In the model the two competing cellular fates adopt different dynamic strategies and shows that there is a lag period of a few days in which the appropriate signalling conditions for PS, namely Nodal/Activin, have to build up population-wise in order to allow cells to become PS. Once this second fate is made available to cells, it becomes the predominant fate (i.e. cells are more prone to become PS than NECT). However, the delay introduced by the signalling mechanism has already allowed the first cells leaving pluripotency to irreversibly commit to the NECT fate. Hence, the interplay between the dynamics of pluripotency exit and the signalling mechanisms allow for a balance between the two opposing fates (Fig. 6C,D).

This situation that we have described in culture is reminiscent of, and provides insights into, the early differentiation in the embryo. The main difference is that in the embryo signalling centres are spatially organized which determines the pattern. It is likely that here there is also a ‘primary’ neural fate [39,75,76] and that PS-promoting signals override this fate. Anterior NECT fates are maintained by the antagonism of PS promoting signals by the secretion of anti-Wnt, anti-Nodal and anti-BMP from the anterior visceral endoderm [16,77,78].

## Materials and Methods

### Routine Cell Culture and differentiation

E14Tg2A, Bra::GFP [32], TNGA [36] and Sox1::GFP [31] mESCs were cultured on gelatin (0.1%) in GMEM (Gibco, UK) supplemented with non-essential amino acids, sodium pyruvate, GlutaMAX™, β-mercaptoethanol, foetal bovine serum and LIF (SL). To obtain serum-free pluripotency conditions, cells were cultured in 2i+LIF; 2i uses an N2B27 base medium (NDiff 227, StemCells Inc., UK) supplemented with 1µM PD0325901, 3µM Chi and LIF. Cell medium was changed daily and cells passaged every other day. For differentiation experiments, cells were plated at a concentration of 4 × 10^3^ cells/cm^2^ in the indicated differentiation medium (day 0). Differentiation medium consisted of an N2B27 base medium supplemented with combinations of Activin (100ng/ml), CHIR99021 (Chi; 3µM), XAV939 (1µM; [79]) SB431542 ([80]; 10µM), BMP4 (1ng/ml) and Dorsomorphin-H1 (0.5µM). Differentiation medium was completely replaced daily to reduce the influence of increased concentrations of secreted factors.

### Flow cytometry and cell sorting

Cells were analyzed for GFP fluorescence using an LSR Fortessa (BD Bioscience) using a 488 laser and emission measured using 530/30 filter, Dapi exclusion was used to determine live cells and measured using 405 laser and emission at 450/50. Data was analyzed using Flowjo software. TNGA cells were sorted according to their GFP fluorescence using a MoFlo sorter (Beckman Coulter) using the same laser and filter sets described above. Cells were collected in SL media, counted and re-plated into the media indicated in the described experiments.

### Immunofluorescence & Confocal Microscopy

Immunofluorescence and image analysis were carried out as described previously [64,71] in 8- well (Ibidi), plastic tissue-culture dishes. Samples were washed in BBS+CaCl_2_ (50 mM BES Sodium Salt, 280 mM NaCl, 1.5 mM Na_2_HPO_4_, 1 mM CaCl_2_ adjusted to pH 6.96 with 1M HCl) and fixed for 15 minutes in 4% paraformaldehyde. Samples were washed and permeablized with BBT (BBS+CaCl_2_ supplemented with 0.5% BSA and 0.5% TritonX-100) before overnight antibody staining, following which, the samples were washed with BBT and incubated for 2 h with the desired fluorescently-conjugated secondary antibody. Prior to imaging, samples were washed with BBS+CaCl_2_ and covered in mounting medium (80% spectrophotometric grade glycerol, 4% w/v n-proply-gallatein in BBS+CaCl_2_). The primary antibodies used were as follows (all at a 1 in 200 dilution): Brachyury (goat; Santa Cruz Biotechnologies, sc17743), Oct3/4 (mouse; Santa Cruz Biotechnologies, sc5279) and Sox2 (rabbit; Millipore, AB5603). Secondary antibodies were from Molecular Probes and used in a 1 in 500 dilution with Hoechst (1 in 1000; Invitrogen). Samples imaged using and LSM700 on a Zeiss Axiovert 200M with a 40x EC Plan-NeoFluar 1.3 NA DIC oil-immersion objective. Hoechst, Alexa488, −568 and −633 were sequentially excited with a 405, 488, 555 and 639 nm diode lasers respectively. Data capture carried out using Zen2010 v6 (Zeiss), image analysis performed using Fiji [81].

### Wide-field Epifluorescence Microscopy

For live imaging, cells were imaged by wide-field microscopy in a humidified CO_2_ incubator (37°C, 5% CO_2_) every 10 min for the required duration using a 20x LD Plan-Neofluar 0.4 NA Ph2 objective with correction collar adjusted for imaging through plastic. All medium was changed daily (see above). An LED, white-light system (Lumencor) was used to excite fluorescent proteins. Emitted light was recorded using an MRm AxioCam (Zeiss) and data recorded with Axiovision (2010) release 4.8.2. Analysis performed using Fiji [81].

### Single Cell qRT-PCR

E14Tg2A wild-type (129/Ola) mES cells were cultured in NDiff N2B27 (StemCells Inc) supplemented with 100U/mL LIF and 10 ng/mL BMP4 (Department of Biochemistry, University of Cambridge). Subsequently, cells were trypsinized and reseeded in N2B27 alone at a density of 10 × 10^3^ cells/cm^2^ to induce differentiation. FACS was used to distribute individual mES cells into the wells of a 96-well plate containing lysis buffer (see below). The transcriptomes of isolated cells were amplified according to a previously published protocol (up to step 30) [82]. The resultant cDNA samples were purified using a PCR purification kit (Qiagen) and diluted 1 in 10 in PCR-grade water.

Diluted cDNA (5µl) was mixed with 9µl Specific Target Amplification mix (STA mix - 7.5µl TaqMan Pre-Amp mastermix (Invitrogen), 1.5µl 10X STA Primer Mix, 0.075µl 0.5M EDTA, pH 8.0). STA Primer Mix is composed of 48 or 96 primer pairs (see supplementary information) diluted in DNA suspension buffer (Teknova) at a concentration of 500nM per primer. Individual amplicons were further amplified using the following thermal cycling protocol: 10 min at 95°C, 20 cycles of 5s at 96°C followed by 4 min at 60°C, 4 min at 60°C. Samples were exonuclease treated and the levels of individual amplicons determined using 48.48 or 96.96 Dynamic Arrays on the BioMark HD platform (Fluidigm) as per the manufacturers instructions (using the EvaGreen based protocol, and the STA primer mix described above). Ct values were extracted automatically using the Biomark Data Collection Software. Melt curves arising from primer-dimer amplification were identified and removed from the dataset manually by comparison with a positive control sample.

### qRT-PCR

RNA was isolated from ∼5 × 10^5^ trypsinized and pelleted mES cells using the RNeasy Mini kit (Qiagen, 74104) according to the manufacturers instructions, and resuspended in 30µL distilled water. RNA was reverse transcribed as follows. RNA samples (1μg in 38μL nuclease free water) were combined with 2μL Oligo-dT anchored primers (Life technologies, 12577-011) and incubated at 80°C for 2 minutes before transferring immediately to ice for 2 minutes. PCR master mix was then added to each sample: 1.5µl dNTPs (Life technologies, 18427-013), 12µl 5X First Strand buffer (Life technologies, 18080-400), 3µl 0.1M DTT (Life technologies, D-1532), 1.5µl RNaseOUT (Life technologies, 10777-019) and 2µl Superscript III Reverse Transcriptase (Life technologies, 18080-400). Thermal cycling was carried out as follows: 25°C 10 minutes, 50°C 30 minutes, 70°C 15 minutes. To remove RNA from the resultant cDNA samples 1µL RNaseH (Life technologies, 18021071) was added and samples were incubated at 37°C for 20 minutes and then at 80°C for 20 minutes. Samples were diluted 1 in 100 before being analyzed for gene expression using the Fluidigm platform (from Specific Target Amplification step using the same primers as for single cell qPCR).

### Mathematical modelling of cell-fate adoption dynamics

The model considers 4 different cellular types: pluripotent cells (*P*), differentiating cells (*D*), cells committed to NECT (*N*) and cells committed to PS (*M*). In the model, we consider initially all cells to be pluripotent. Cells then are allowed to differentiate at a certain rate *λ_D_*. Once each cell abandons the pluripotent state, it rapidly and irreversibly adopts a final fate: either NECT or PS. Cells are assumed to acquire either fate at a particular rate: *λ_N_* for NECT and *λ_M_* for PS. In addition to these cellular types and the corresponding transitions, we also consider a signal (*s*) that builds up as cells loose pluripotency. Once the signal crosses a certain threshold, it biases the rates of fate adoption. In particular, we consider that the rate of PS conversion is negligible in the absence of signal (*λ_M_ << λ_N_*) and becomes the much larger (*λ_M_ >> λ_N_*) once the signal reaches the threshold. In this simple model we disregard both cellular death and cell division, assuming that both processes are effectively balanced within all cellular fates. The model equations are described in Fig. 6 Supplementary Data A. These define a simple linear ODE system with a piecewise constant rate *λ_M_*, (which depends on the levels of accumulated signal). Thus a solution can be built up by gluing the analytical solutions of each range of constant *λ_M_* (Fig. 6 Supplementary Data B). Two of the three model parameters, *λ_D_* and *λ_N_*, have been fitted to the experimental data of N2B27 differentiation by assuming that no differentiation to PS occurs, i.e., *λ_M_*=0 (Fig. 6 Supplementary Data C,D). With the obtained parameter values, and assuming a switch in the dynamic regime (*λ_M_ >> λ_N_*) in the presence of Activin and Chiron, the model reproduces to a great extent the experimental pulse-chase results (cf. Fig. 1D and Fig. 6 Supplementary Data E).

## Acknowledgements

The authors thank the Wellcome Trust (JT) and the ERC (AMA, DAT, PH and PR) for funding.

